# Human AdV-20-42-42, a promising novel adenoviral vector for gene therapy and vaccine product development

**DOI:** 10.1101/2021.03.11.435055

**Authors:** Mónika Z. Ballmann, Svjetlana Raus, Ruben Engelhart, Győző L. Kaján, Chantal van der Zalm, Tibor Papp, Lijo John, Selina Khan, Jerome Custers, Wilfried A.M. Bakker, Hilde M. van der Schaar, Niklas Arnberg, Angelique A.C. Lemckert, Menzo Havenga, Andrew H. Baker

**Affiliations:** Batavia Biosciences B.V., Leiden, The Netherlands; Centre for Cardiovascular Sciences, University of Edinburgh, Little France Crescent, Edinburgh, UK; Department of Clinical Microbiology, Division of Virology, Umeå University, Sweden; Janssen Vaccines and Prevention B.V., Leiden, The Netherlands

**Keywords:** Novel adenovirus serotype, Expression vector, Low seroprevalence, Cell and tissue transduction, Potent T-cell responses

## Abstract

Pre-existing immune responses towards adenoviral vector limit the use of a vector based on particular serotypes and its clinical applicability for gene therapy and/or vaccination. Therefore, there is a significant interest to vectorize novel adenoviral types that have low seroprevalence in the human population. Here, we describe the discovery and vectorization of a chimeric human adenovirus, which we call HAdV-20-42-42. Full genome sequencing revealed that this virus is closely related to human serotype 42, except for the penton-base which is derived from serotype 20. The HAdV-20-42-42 vector could be propagated stably to high titers on existing E1-complementing packaging cell lines. Receptor binding studies revealed that the vector utilized both CAR and CD46 as receptors for cell entry. Furthermore, the HAdV-20-42-42 vector was potent in transducing human and murine cardiovascular cells and tissues, irrespective of the presence of blood coagulation factor X. In addition, the vector did not sequester in the liver upon intravenous administration in rodents. Finally, we demonstrate that potent T-cell responses against vector-delivered antigens could be induced upon vaccination. In summary, from the data obtained we conclude that HAdV-20-42-42 provides a valuable addition to the portfolio of adenoviral vectors available to develop safe and efficacious products in the fields of gene therapy and vaccination.

**IMPORTANCE:** Adenoviral vectors are currently under investigation for a broad range of therapeutic indications in diverse fields, such as oncology and gene therapy, as well as for vaccination both for human and veterinary use. A wealth of data shows that pre-existing immune responses may limit the use of a vector. Particularly in the current climate of global pandemic, there is a need to expand the toolbox with novel adenoviral vectors for vaccine development. Our data demonstrates that we have successfully vectorized a novel adenovirus serotype with low seroprevalence. The cell transduction data and antigen-specific immune responses induced *in vivo* demonstrate that this vector is highly promising for the development of gene therapy and vaccine products.

## INTRODUCTION

Adenoviral vectors have been studied for decades as they hold great promise as tools to develop safe and effective gene therapy and vaccine products. As such, there are dozens of therapeutic applications being pursued utilizing adenoviral vectors. As it has been amply demonstrated that host immune responses limit the repeated use of a particular vector (1-4) there is a constant demand to identify new adenovirus (AdV) serotypes with low seroprevalence, and alternate tropism. Thus, investigations into the *in vitro* and *in vivo* biology of less prevalent adenovirus may advance the clinical use of alternative Ad-based platforms.

To date 57 human adenovirus (HAdV) serotypes have been described and are sub-grouped in 7 species (HAdV A-G), of which HAdV-D is the largest. It has been well-documented that the members of the different species are associated with diverse clinical symptoms, including gastroenteritis (HAdV-F and -G), respiratory disease (HAdV-B, C, and E), or conjunctivitis and/or keratitis (HAdV-B and -D). HAdV-induced symptoms can either be self-limiting and cleared by a host within days to weeks, but persistent infection of HAdV-C can last for months. Serotype classification is historically based on the unique neutralization profile of an adenovirus, i.e. serum specifically raised against one serotype does not cross-neutralize other adenovirus serotypes, and a hemagglutination profile. Besides the 57 acknowledged serotypes, many hybrid adenoviruses have been discovered and as these so-called chimeras can be neutralized by parental serum they do not qualify as distinct serotypes. Most likely such hybrids originate from homologous recombination events between two or more viruses replicating simultaneously in a host cell during co-infection (5-8).

In the present study we describe the generation of a novel replication-incompetent vector based on a natural hybrid that we named HAdV-20-42-42. The recombinant HAdV-20-42-42 vector was used for characterization of its seroprevalence, receptor usage, and tropism. In addition, we explored the utility of the vector as a potential tool to develop gene therapy and vaccine products. The data obtained and described here warrant further studies into the utilization of the HAdV-20-42-42 vector to develop vaccines and cardiovascular intervention strategies.

## RESULTS

### Identification of HAdV-20-42-42, a natural chimera

In order to identify possible new vector candidates, e.g. new and/or rare human adenovirus types, 281 human adenovirus strains isolated from patients in Sweden between 1978 and 2010 were screened previously (9). From these samples, the hexon, the penton base, and the polymerase genes were amplified and sequenced to allow for identification of new adenovirus types or possible recombinants. Strains with interesting genotypes were propagated on A549 cells, after which the complete viral genome was sequenced using Next-Generation Sequencing and annotated based on HAdV reference strains. Phylogenetic analyses were conducted based on the complete genome sequence and amino acid translations of the hexon, penton base, and the fiber knob.

During the screening process, strain 212 was pinpointed and analyzed further. The complete genome sequence of this strain was 35,187 bp long with a GC-content of 57.0%. A typical HAdV-D genome layout was observed with 37 protein coding sequences and two virus-associated RNAs (Figure 1A), pointing to a recombination event of two types that resulted in a hybrid. The strain is closest related to HAdV-42 (species *Human mastadenovirus D*) in most phylogenetic analyses except for the penton base, which has the highest sequence identity with HAdV-20 (Figure 1B). Thus, the genomic composition of strain 212 was determined as HAdV-20-42-42 concerning the sequence of the penton base, the hexon, and the fiber knob.

**Figure 1A.**
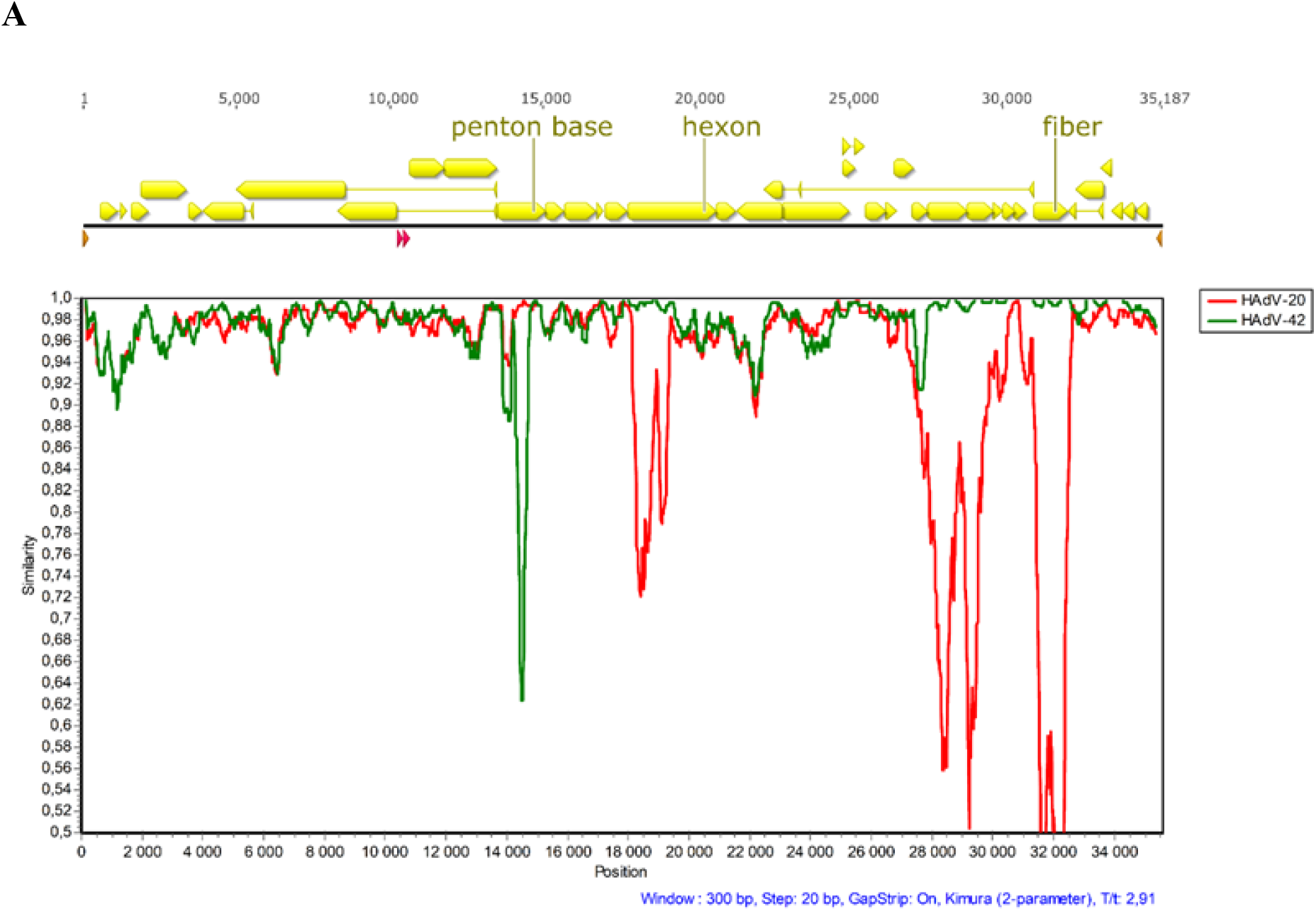
Identification of HAdV-20-42-42, a natural chimera. **A)** Yellow arrows in the genome map represent protein coding sequences, red arrows represent virus associated RNAs and brown arrows represent the inverted terminal repeats. In the SimPlot analysis, sequence identities to human adenovirus 20 and −42 are represented by red and green plots, respectively.

**Figure 1B.**
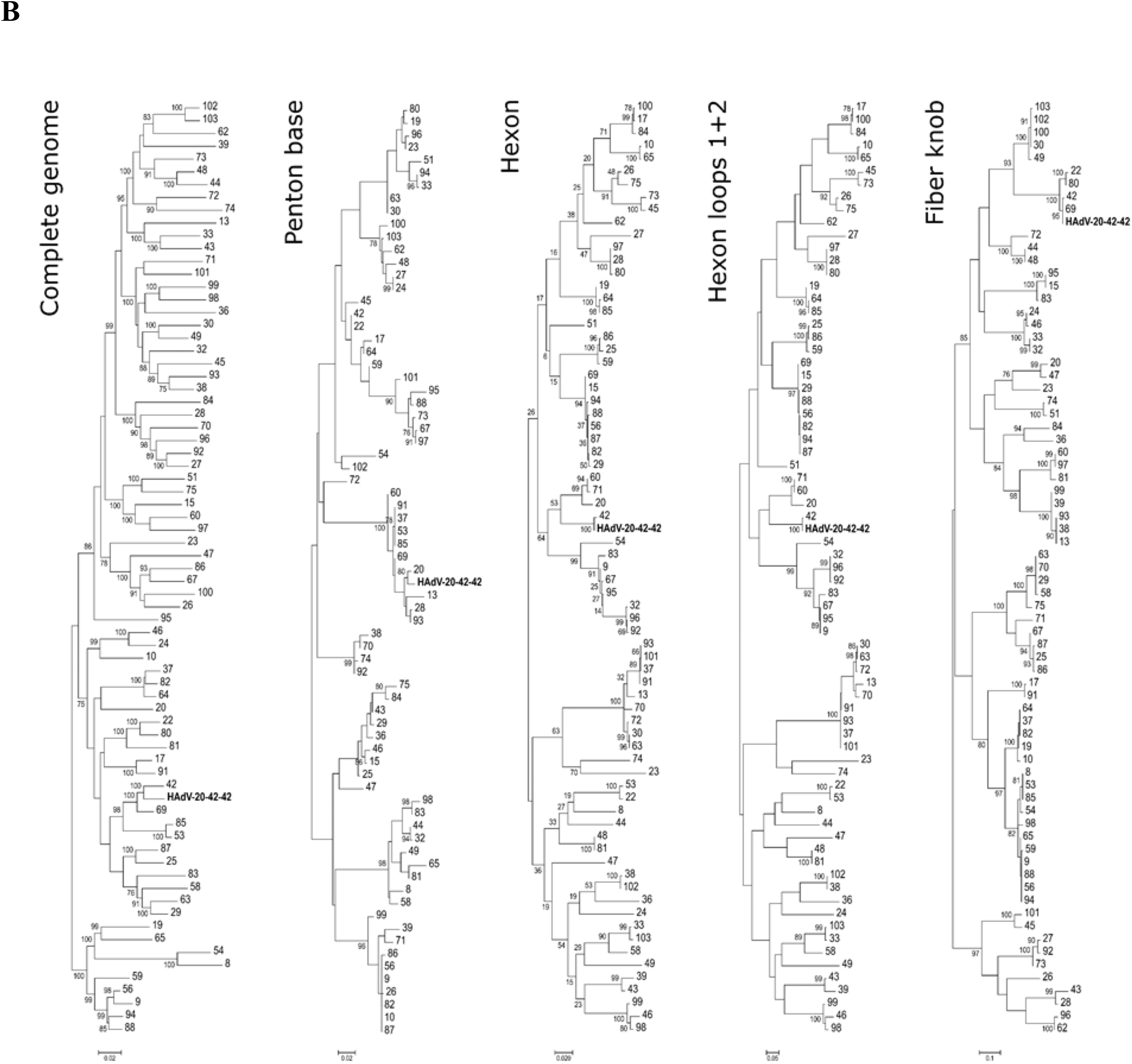
Identification of HAdV-20-42-42, a natural chimera. **B)** Phylogenetic analysis of strain 212 (Umu009) based on the complete genome sequence and derived amino acid sequences of the hexon, the penton base, the hexon loop 1 and the fiber knob. *Human mastadenovirus D* reference strains are represented by their serotype- or genotype numbers.

### HAdV-20-42-42 shows low seroprevalence in human subjects

High levels of pre-existing anti-vector humoral immunity in vaccine target populations may hamper potential use of a novel adenoviral vector as an efficacious vaccine platform, such as found for HAdV-5-based vectors (10-12). We investigated the levels of pre-existing neutralizing antibodies against HAdV-20-42-42 using a panel of serum samples (*n*=103) taken from a cohort of healthy >50-year-old US citizens. In line with previous findings (13), ∼60 % of the serum samples exhibited high levels of neutralizing activity (effective at dilutions >1 : 200) against HAdV-5 (Fig. 2). For HAdV-35 and HAdV-20-42-42 on the other hand only ∼15 % of serum samples neutralized viral activity at dilutions >200 (Fig. 2). These data indicate that the seroprevalence of HAdV-20-42-42 in this cohort was low with antibody levels comparable to that of the rare serotype HAdV-35 (species HAdV-B).

**Figure 2.**
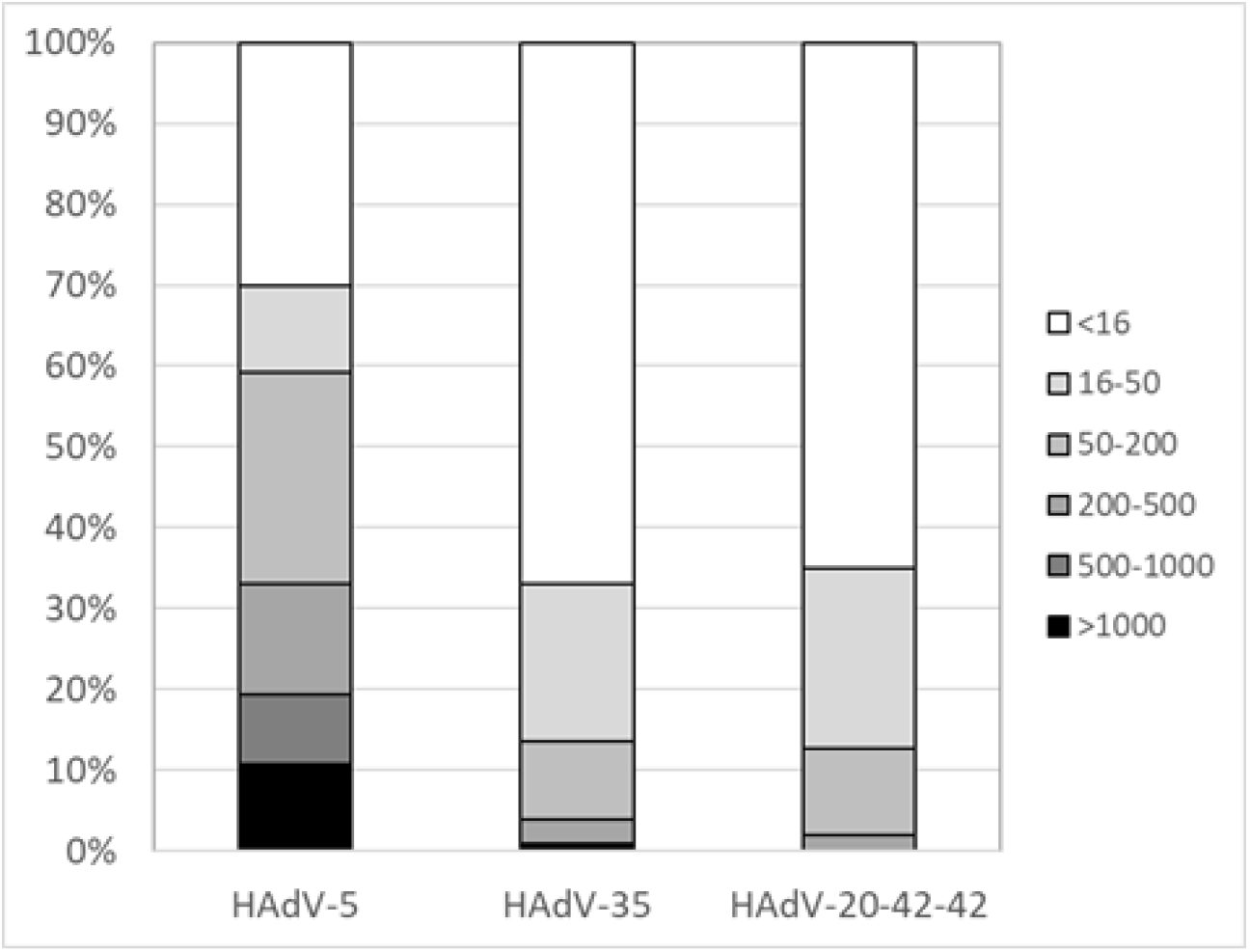
HAdV-20-42-42 shows low seroprevalence in studies with human subjects. HAdV-5, HAdV-35 and HAdV-20-42-42 seroneutralization by a cohort of healthy >50-year-old US citizens (*n*=103 individual serum samples). The neutralization titers were arbitrarily divided into the following categories: <16 (no neutralization), 16 to 300, 300 to 1,000, 1000 to 4000 and >4000.

### Vectorization of HAdV-20-42-42

In order to study the therapeutic applicability of HAdV-20-42-42, we first generated replication-incompetent vectors expressing reporter genes *β*-galactosidase (LacZ), luciferase (Luc) or Enhanced Green Fluorescent Protein (EGFP). To do so, engineered HAdV-20-42-42 genomic DNA sequences were cloned into three plasmids, called the adaptor plasmid, the intermediate plasmid, and the right-end plasmid, with overlapping regions to allow for homologous recombination events (see Figure 3A). To produce replication-incompetent reporter viruses, the E1 region of HAdV-20-42-42 was replaced with an expression cassette and a reporter gene in the adaptor plasmid. In the right-end plasmid, the E3 region was deleted to create capacity for insertion of larger transgenes. To enhance the growth in a standard producer cell line (i.e. HAdV-5 E1-complementing HEK293 cells), the native E4 ORF6/7 region was exchanged with that of HAdV-5 (13-15).

**Figure 3.**
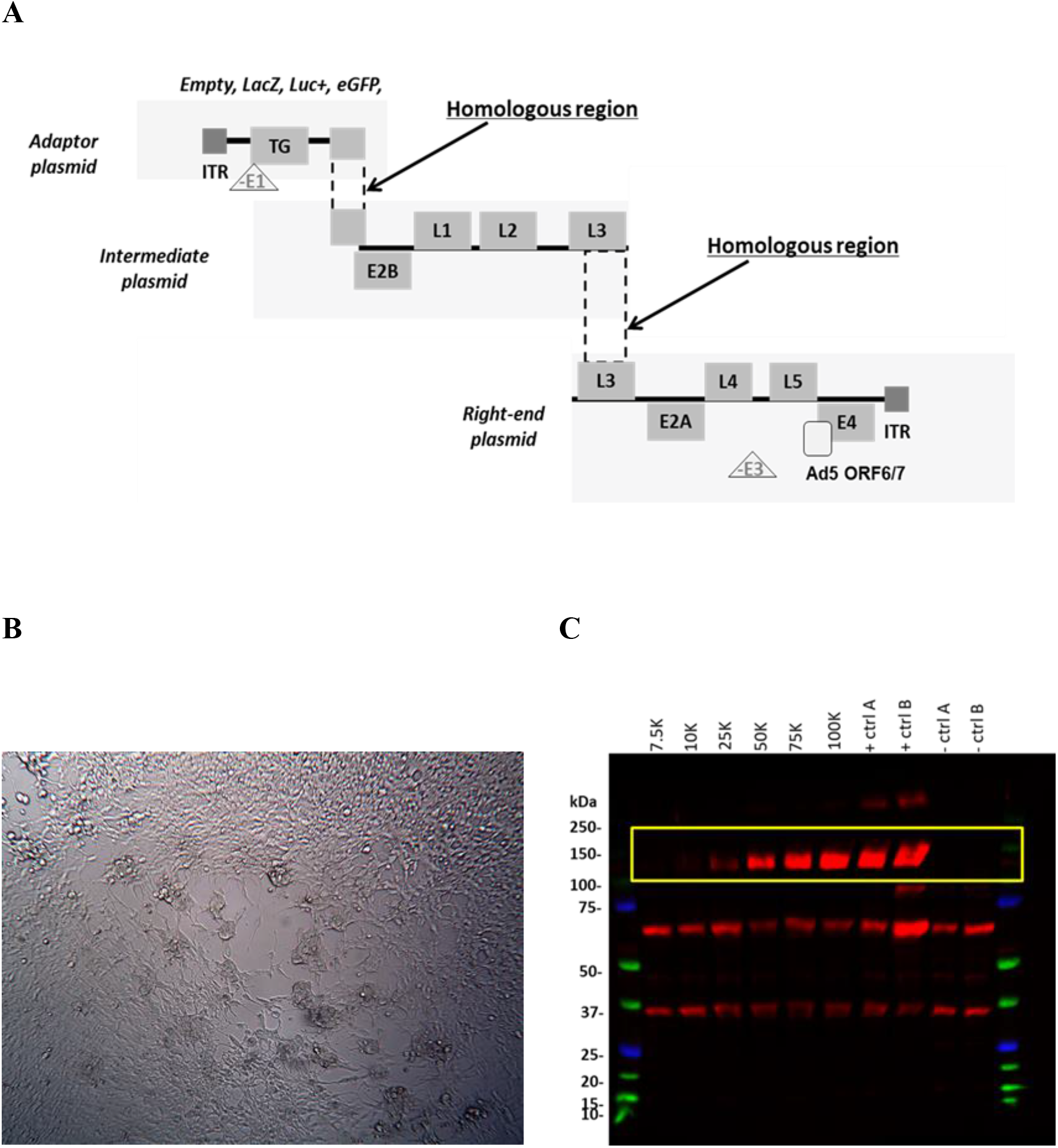
HAdV-20-42-42 vector generation and characterization. **A)** Schematic representation of the replication-incompetent HAdV-20-42-42 recombinant viral vector construction strategy with a three-plasmid system (adaptor, intermediate and right-end plasmid). Overlapping regions allowed homologous recombination events between the HAdV-20-42-42 sequences in HEK293 production cell line. E and L represent the “early” and “late” genes of the adenoviral genome, ITRs are the inverted terminal repeats at the 5’ and 3’ends. **B)** CPE development in HEK293 cells upon transfection of three HAdV-20-42-42 plasmids. **C)** Verification of transgene (LacZ) expression. A549 cells were infected with HAdV-20-42-42-LacZ at various MOIs (indicated as vp per well). At 3 dpi cells were lysed and subjected to Western Blot analysis with an antibody against LacZ. The LacZ-specific bands are marked by a yellow rectangle. Lysates of a previous HAdV-20-42-42-LacZ infection were used as controls (+ ctrl A and B), while lysates of HAdV-20-42-42-GFP infected cells (-ctrl A) or uninfected cells (-ctrl B) were used as negative controls.

Reconstruction of the full-length recombinant HAdV-20-42-42 vector encoding the various reporter genes was achieved by homologous recombination via co-transfection of the three plasmids into HEK293 cells. After co-transfection, the HEK293 cells were subjected to a freeze-thaw cycle to release intracellular particles. After removal of cell debris by centrifugation, the supernatant was used for a reinfection of fresh HEK293 cells. At 3 days post-re-infection of the fresh HEK293 cells cytopathic effect (CPE) was observed (Figure 3B) and viral progeny were successfully propagated to high titers and purified with CsCl density gradients. The reporter gene expression was detected successfully in infected cells (Figure 3C).

### Characterization of HAdV-20-42-42 receptor usage

Well-studied adenovirus gene therapy or vaccination vectors (e.g. HAdV-5, HAdV-26 and HAdV-35) bind their fibers to CD46, coxsackie and adenovirus receptor (CAR) or desmoglein 2 (DSG2) as high affinity receptors, while their penton bases interact with αν-integrins as co-receptors for internalization (16). HAdV-20-42-42 clusters to a small group of HAdV-Ds, which have been previously shown to utilize subunits of sialic acid as primary receptors for cellular attachment and/or entry. To investigate the receptor usage, we determined the transduction capacity of HAdV-20-42-42-Luc in various cell lines, expressing or lacking CAR, sialic acid-containing glycans, DSG2, or CD46 isoforms, by measuring the luciferase levels after infection. Similar to HAdV-5, HAdV-20-42-42 efficiently transduced CHO-CAR cells, but was also able to transduce Chinese hamster ovary (CHO) cells lacking the CAR receptor albeit at a much lower efficiency (Fig. 4A), suggesting that it may also utilize other receptors.

**Figure 4.**
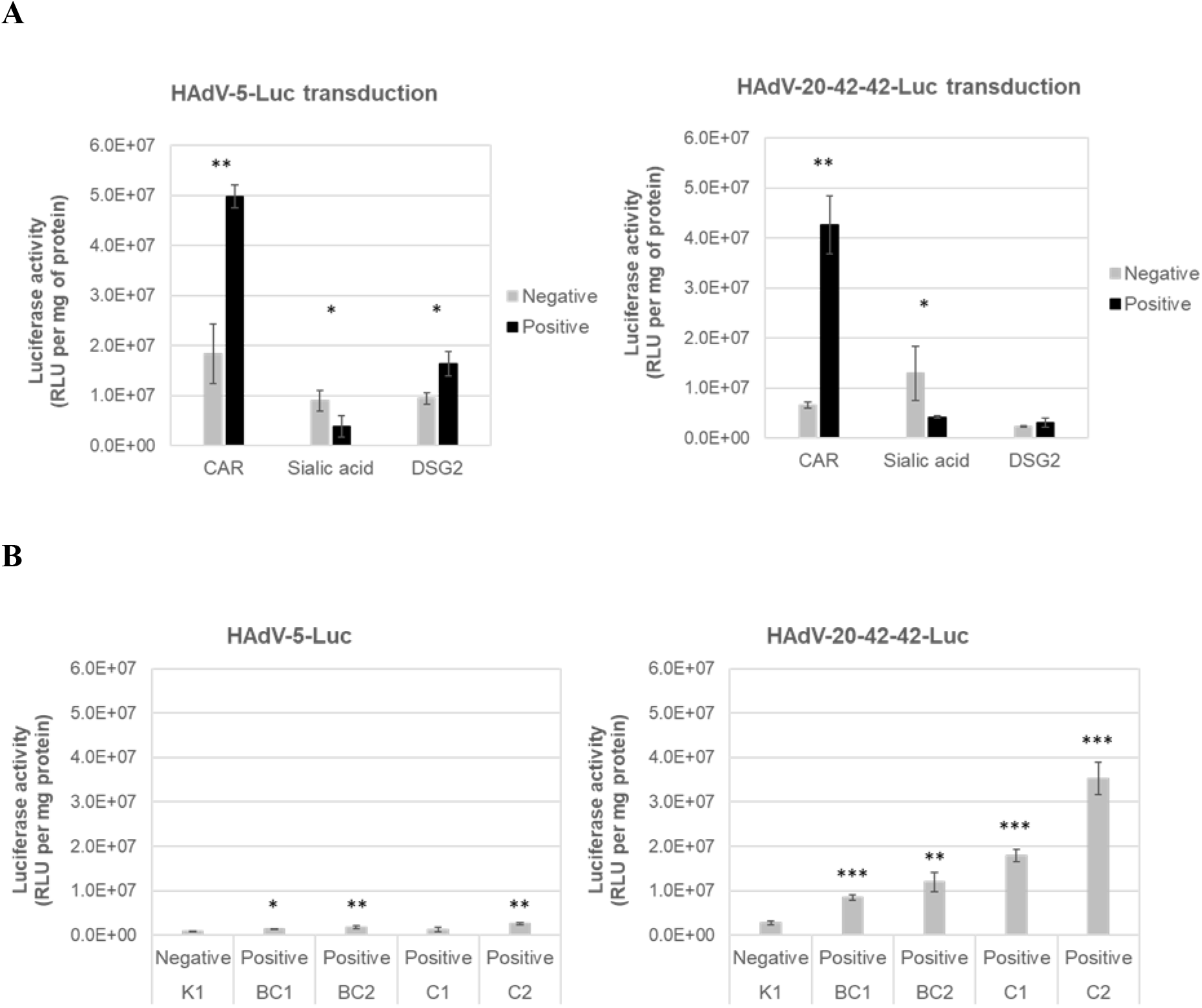
Receptor usage of HAdV-20-42-42. **A)** Cells expressing (positive, dark gray bars) or lacking (negative, light gray bars) CAR, sialic acid-containing glycans or DSG2 receptors were infected with HAdV-20-42-42-Luc or HAdV-5-Luc as a control vector. **B)** CHO cells lacking CD46 (K1) or expressing various CD46 isoforms (BC1, BC2, C1, or C2) were infected with HAdV-20-42-42-Luc or HAdV-5-Luc as a control vector. **A, B)** One day post-infection, cells were lysed to determine intracellular luciferase activity. Luciferase activity is presented as relative light units (RLU) per milligram (mg) protein. All results represent averaged data from several times performed experiments, with four replicates for each condition. Error bars are presented as standard error of the mean (SEM). Two-sample, two-tailed Student’s t-tests, p<0.05 was considered statistically significant (p<0.05*, p<0.01* *). Statistical significance was calculated for positive versus negative cells (**A**) or in comparison to the negative cell line K1 (**B**).

Although being exploited by other HAdV-D members for infection, the presence of sialic acid on cells did not increase the transduction capacity of HAdV-20-42-42. Levels of DSG2 in monocytes also did not affect the luciferase expression of HAdV-20-42-42 (Fig. 4A). Next, the transduction efficiency was evaluated in CHO cells expressing various isoforms of CD46 (Fig. 4B). CHO-K1 cells, which lack CD46, were poorly transduced by HAdV-20-42-42. While HAdV-5 interacted with all CD46 isoforms as well as with CHO-K1 cells, HAdV-20-42-42 preferentially used the C2 receptor isoform of CD46 for infection (Fig. 4B). Together, these findings indicate that the novel adenoviral vector HAdV-20-42-42 is able to bind to both CAR and CD46 receptors, primarily the C2 isoform. As these receptors are present in many cell types, this warrants a broad use of the novel adenoviral vector HAdV-20-42-42 in gene therapy.

### HAdV-20-42-42 vector interactions with serum and coagulation factors

The binding of many HAdV types to human coagulation blood factor X (FX) significantly affects the transduction *in vitro* and the tropism *in vivo* following intravenous (i.v.) administration (10, 17, 18). We investigated whether the presence of FX impacts the transduction efficiency of HAdV-20-42-42. Physiological concentrations of FX were incubated with cells prior to the addition of HAdV-5, −35, or HAdV-20-42-42 luciferase vectors, and intracellular luciferase levels were measured 2 days after transduction. While HAdV-35 transduction was only marginally affected by the addition of FX, the luciferase levels of HAdV-5 and in particular of HAdV-20-42-42 were substantially increased in the presence of FX (Fig. 5A). These data show a notably higher transduction potential of HAdV-20-42-42 over HAdV-5 and HAdV-35 in human saphenous vein endothelial cells (HSVEC) in the presence of FX.. HSVEC were infected with HAdV-20-42-42-LacZ and HAdV-5-LacZ as a control. At all doses tested, the percentage of LacZ-positive cells was higher for HAdV-20-42-42 compared to HAdV-5 (Fig. 5B). These data show that HAdV-20-42-42 was capable of efficiently transducing vascular cells in the presence of FX.

**Figure 5.**
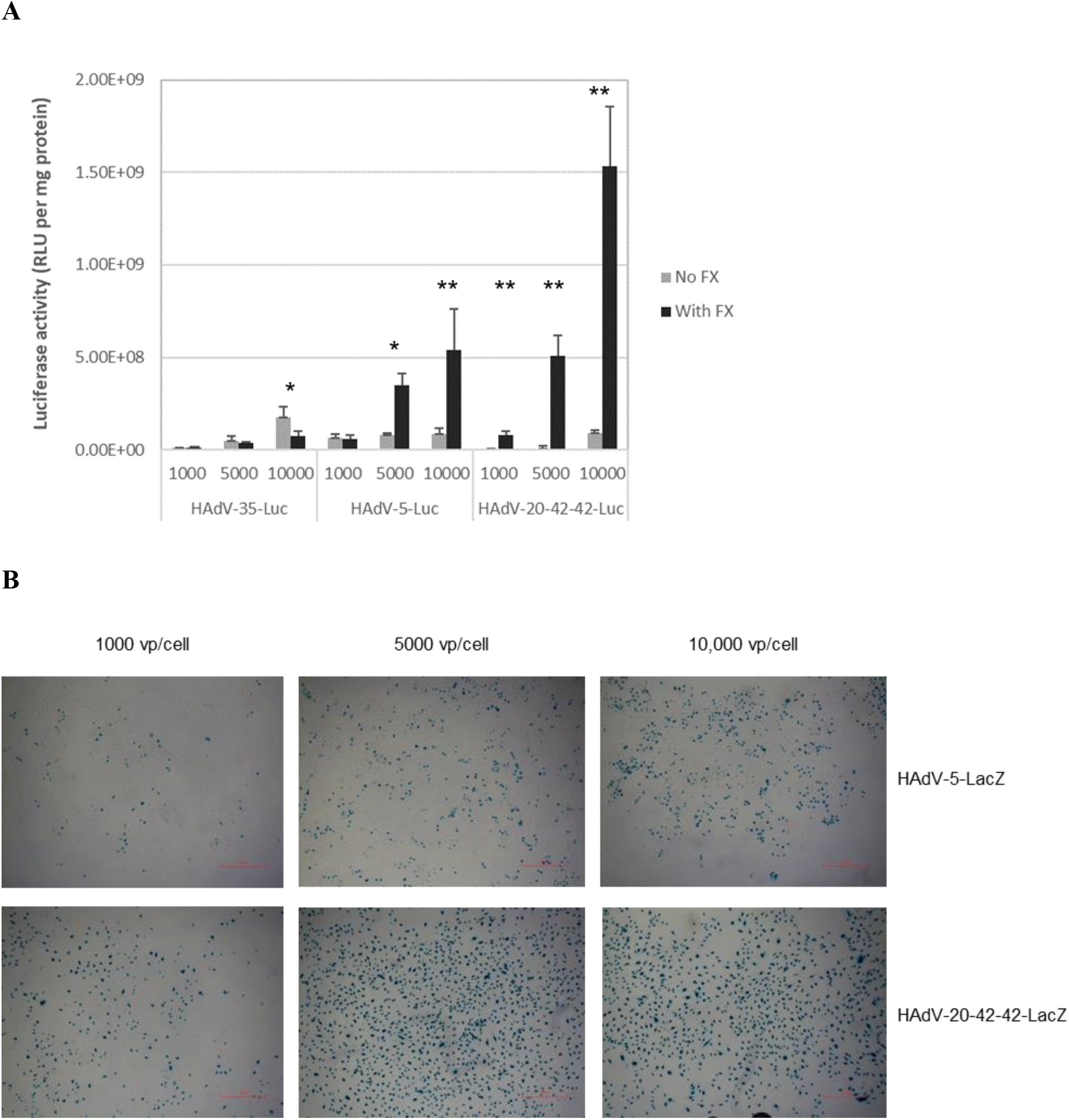
Transduction efficiency of HAdV-20-42-42 in vascular cells. HAdV-20-42-42-Luc and -LacZ vectors were tested for transduction capacity in HSVEC (human saphenous vein endothelial cells). **A)** HSVEC were infected with HAdV-20-42-42-Luc or the control vectors HAdV-35-Luc and HAdV-5-Luc at various doses (1000, 5000, or 10000 vp/cell) for 3 hours. Where indicated (dark gray bars), the vectors were incubated for 30 min at 37°C with a physiological concentration of 10 µg/ml blood coagulation factor FX prior to addition to the cells. After 2 days, cells were lysed to measure intracellular luciferase activity, which is presented as relative light units (RLU) per milligram (mg) protein. Bars represent the means plus standard errors of the means (SEM) (error bars) for quadruplicate values. Two-sample, two-tailed Student’s t-tests, p<0.05 was considered statistically significant (p<0.05*, p<0.01* *). All results represent averaged data from several experiments, with four replicates for each condition. **B)** HSVEC were infected with various doses of HAdV-20-42-42-LacZ or HAdV-5-LacZ in the presence of FX. After 2 days, cells were stained for LacZ expression.

### HAdV-20-42-42 biodistribution following systemic delivery *in vivo*

To characterize the HAdV-20-42-42 vectors *in vivo* following intravenous administration, we evaluated the biodistribution patterns of HAdV-20-42-42-Luc using a previously described mouse model (19, 20). Animals inoculated with vehicle (PBS) or HAdV-5-Luc were included as negative and positive control groups, respectively. Pretreatment with clodronate liposomes (CL+) was performed on half of the animals in order to deplete macrophages, thereby reducing possible sequestration of the adenovirus vector to the liver and allowing a more efficient evaluation of the biodistribution at the whole organism level. Two days after vector administration, the animals were imaged to visualize luciferase levels (Fig. 6A). Subsequently, the mice were sacrificed and several organs were collected for adenoviral DNA quantification (Fig. 6B).

**Figure 6.**
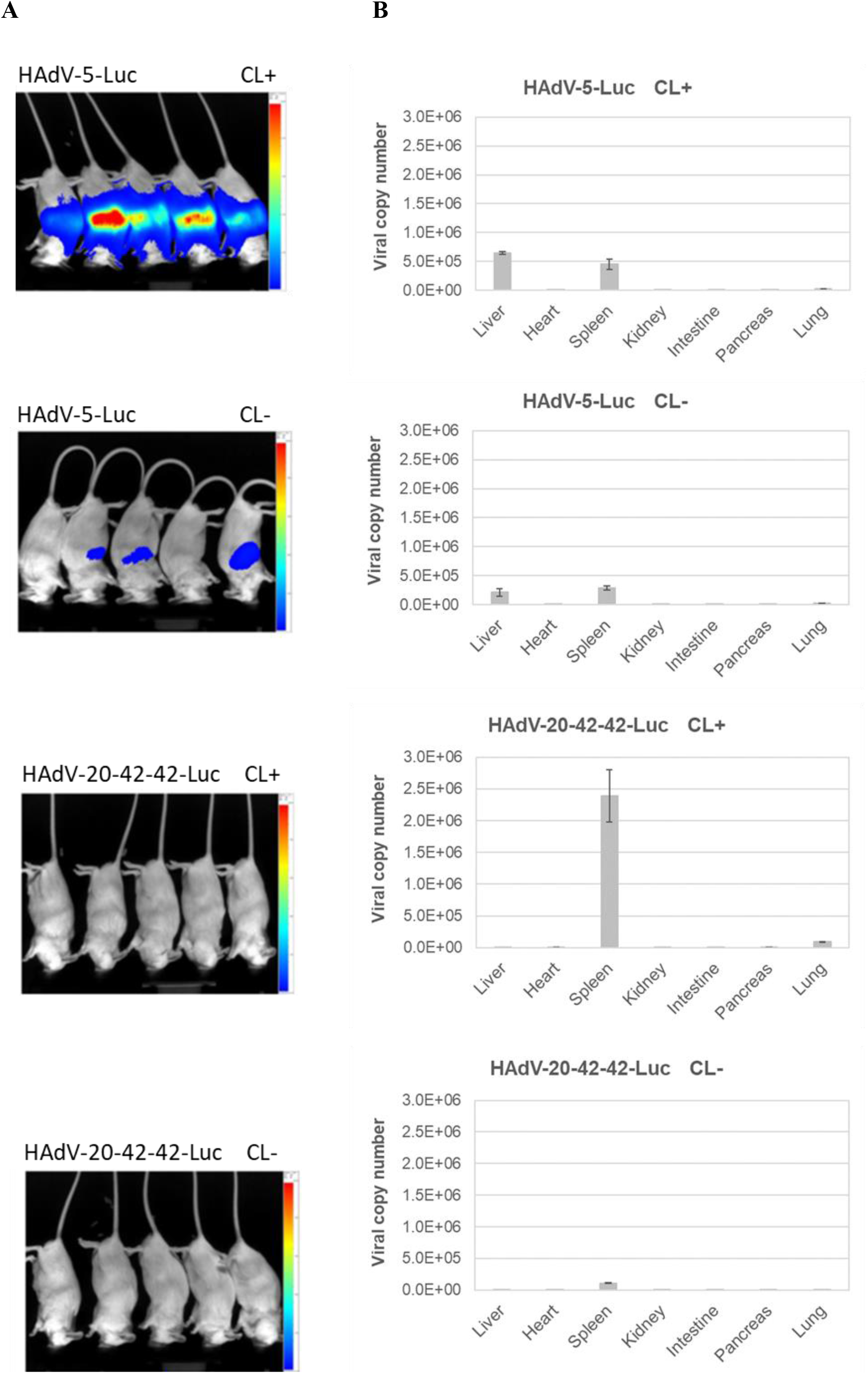
Biodistribution profile of HAdV-20-42-42 shows mainly spleen tropism. Mice were pre-treated with clodronate (CL+) or untreated (CL-) and injected intravenously with HAdV-5-Luc or HAdV-20-42-42-Luc vectors. **A)** At 48 hours post virus delivery, luciferin was injected to the mice and luciferase activity was imaged with the method IVIS Spectrum, which ranged from low activity (shown in blue) to high activity (shown in red) levels. **B)** After imaging, animals were sacrificed and organs were collected to determine adenoviral genome copy numbers with qPCR. Data represent viral copy number per 100 ng of total DNA. Bars represent the means plus standard errors of the means (SEM).

In the control group of animals without CL pretreatment, HAdV-5-Luc was mainly distributed in liver and spleen at the levels of ∼2.5 × 10^5^ genome copy numbers per 100 ng total DNA, while in the group pre-treated with CL the liver and spleen distribution was higher, closer to ∼5 × 10^5^ genome copy numbers (Fig. 6). HAdV-20-42-42 on the other hand appeared to have only a spleen tropism, since luciferase activity was not detected in other organs. As expected, the total DNA copy number was significantly higher when CL was added (∼2.5 × 106) in comparison to the group without CL pretreatment (1 × 10^5^).

Collectively, these data indicate that HAdV-20-42-42 has a good safety profile with only spleen tropism found in the studies, while the adenoviral DNA copies in other organs tested were poorly detectable.

### HAdV-20-42-42 as a candidate vaccine vector

The potential of HAdV-20-42-42 as a vaccine vector candidate was assessed for its ability to induce cellular immune responses against a model antigen (Luciferase, Luc) in Balb/c mice after intramuscular immunization. The vector was compared side-by-side with a benchmark vector based on HAdV-26, which has undergone clinical trials for HIV, Ebola, and recently for SARS-coronavirus-2 (21-24). Mice were immunized intramuscularly with two different doses of E1- and E3-deleted HAdV-26 vector expressing luciferase (HAdV-26-Luc) or HAdV-20-42-42-Luc. Mice were sacrificed and sampled for serum and splenocytes two weeks after the prime immunization. Cellular immune responses against the vector-encoded antigen was evaluated by Luc specific-IFN-γ ELISPOT assay. To this end, splenocytes sampled from immunized mice were stimulated overnight with a 15mer overlapping Luc peptide pool. The antigen specific immune responses were determined by measuring the relative number of IFN-γ-secreting cells (Fig. 7A). As expected, no Luc specific responses were detected against the empty adenovectors lacking luciferase (HAdV-26.E). Furthermore, the results show that the cellular immune responses induced by HAdV-20-42-42 were comparable to the response seen for HAdV-26 at the highest immunization dose (10^10^ VP).

**Figure 7.**
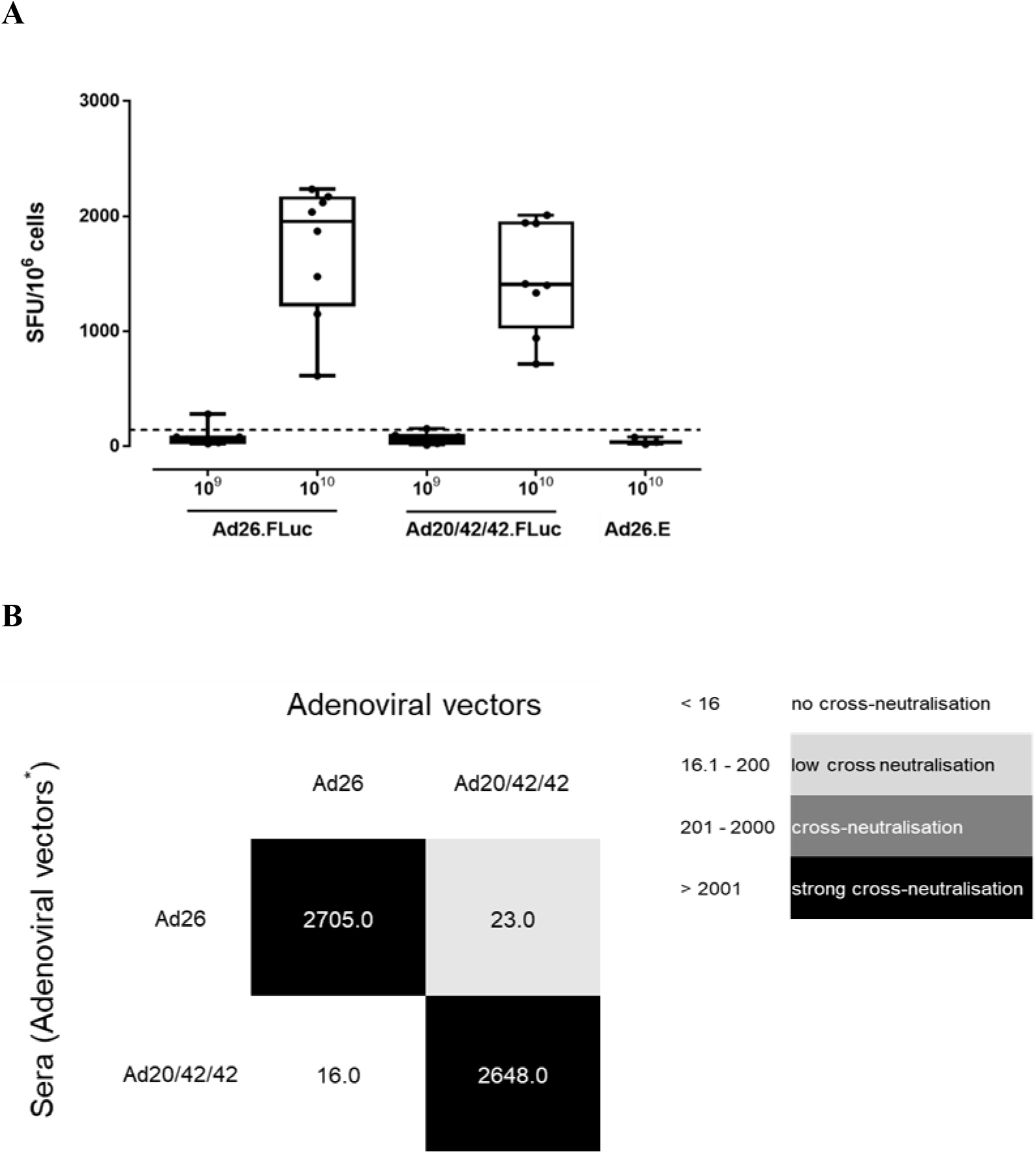
Candidate vector HAdV-20-42-42 elicits strong immune response in mice. **A)** Cellular immune response in mice. Balb/C mice were immunized by intramuscular injection with HAdV-26-Luc, HAdV-20-42-42-Luc, or with HAdV-26-E that lacks a transgene. Two vector doses, i.e. 10^9^ and 10^10^ viral particles per mouse, were administered and animals were sacrificed two weeks post immunization and sampled for serum and splenocytes. Cellular immune responses against the vector-encoded antigen was evaluated by Luc specific-IFN-γ ELISPOT assay. To this end, splenocytes were stimulated overnight with a 15mer overlapping Luc peptide pool. The antigen specific immune responses were determined by measuring the relative number of IFN-γ-secreting cells. shown as Spot Forming Units (SFU) per million cells. Each dot represents a mouse, the bar indicates the geometric mean and the dotted line is the 95^th^ percentile based on the medium control samples. **B)** Cross-neutralization between HAdV-20-42-42 and HAdV-26. Mice antisera against hAd20V-42-42 and HAdV-26 were cross-tested against both vectors in an adenovirus neutralization assay. Starting from a 1:16 dilution, the sera were 2-fold serially diluted, then pre-mixed with the adenoviral vectors expressing luciferase, and subsequently incubated overnight with A549 cells at MOI 500 virus particles. Luciferase activity levels in infected cell lysates measured 24 hours post-infection represented vector infection efficiencies. The neutralization titers were arbitrarily divided into the categories shown in the legend on the right.

For their potential utility as new adenoviral vaccine vectors, the novel HAdV-20-42-42 adenoviral vector would preferably be serologically distinct from existing adenoviral vectors currently in development as vaccine vectors, such as HAdV-26. Therefore, cross-neutralization tests were performed between the novel HAdV-20-42-42 adenoviral vector and HAdV-26. To this end, mice antisera raised against these vectors during the immunization study described above were cross-tested against both vectors in an adenovirus neutralization assay. The adenovirus neutralization assay was carried out as described previously (25). Briefly, starting from a 1:16 dilution, the sera were 2-fold serially diluted, then pre-mixed with the adenoviral vectors expressing luciferase (Luc), and subsequently incubated overnight with A549 cells at a multiplicity of infection (MOI) of 500. Luciferase activity levels in infected cell lysates measured 24 hours post-infection represented vector infection efficiencies. Neutralization titers against a given vector were defined as the highest serum dilution capable of giving a 90% reduction of vector infection efficiency. The neutralization titers were arbitrarily divided into the following categories: <16 (no neutralization), 16.1 to 200 (low cross-neutralization), 201 to 2,000 (cross-neutralization), and >2,001 (strong cross-neutralization).

The results show no major cross-neutralization between the vectors tested (Fig. 7B), but a low, one-way cross-neutralization was observed with HAdV-26 antiserum displaying a neutralization titer against HAdV-20-42-42 of 23. Thus, the new adenoviral vector HAdV-20-42-42 displayed low, if any, cross-neutralization with the human adenoviral vector HAdV-26.

## DISCUSSION

We present the first report on the generation of a replication-incompetent HAdV-20-42-42 vector and present data on initial *in vitro* and *in vivo* characterization. Our serum neutralization studies using wildtype HAdV-20-42-42 virus demonstrated that this virus displayed low seroprevalence in a random set of sera derived from healthy US subjects. Low seroprevalence has been an important criterium to select a virus for vector development, as it has been amply demonstrated that the transduction capability of the vector can be hampered in the presence of a high titer of neutralizing antibodies (26).

Based on these initial data we set out to generate a vector system based on HAdV-20-42-42. We created a flexible three plasmid system to support HAdV-20-42-42 vector generation, allowing for the convenient insertion of transgenes into a multiple cloning site. The wildtype HAdV-20-42-42 E4ORF6 region was replaced with that of Ad5, a technique that we previously adopted for the generation of other species HAdV-D viruses including HAdV-26, HAdV-48, HAdV-49 and HAdV-56 vectors. This replacement allowed for their successful production in Ad5 E1-complementing cells such as HEK293 and PER.C6^®^ (13, 27, 28) and enabled us to manufacture high quality, high titer HAdV-20-42-42 vectors carrying diverse inserts, which paves the road for large scale clinical productions.

Previous studies have demonstrated that HAdV-D strains can employ CAR, CD46, sialic acid-containing glycans and αv-integrins as entry receptors (16). In accordance with this, we demonstrate here that HAdV-20-42-42 utilizes CAR and CD46 as receptors. Similar to HAdV-26 and HAdV-48, this virus interacts with blood coagulation factors (10). Together with data demonstrating that HAdV-20-42-42 bypasses the liver, resulting in the absence of hepatoxicity, as well as the high transduction efficiency of cardiovascular cells in vitro, these findings suggest that this vector is potentially well suited to develop cardiovascular gene therapy approaches although further in vivo studies are required to develop this concept in more detail. In addition, the data obtained from our vaccination experiments suggest that this vector is capable of eliciting potent insert specific T-cell responses at similar levels as compared to a HAdV-26 vector. The HAdV-26 vector platform has recently been successfully used to develop a potent and safe vaccine against Ebola and this vaccine has been approved for human use by European regulatory authorities. In addition, this vector is being tested for the development of preventive vaccines against HIV, RSV and more recently SARS-CoV-2 (21, 22, 29). In preclinical tests, HAdV-26 used alone was demonstrated to induce robust humoral and cellular immunity, plus it has been well tolerated in humans while eliciting target specific immunity in phase I-III trials. We were interested to test cross-neutralization between both D viruses to assess whether subsequent use of these vectors would be a possibility. Our data demonstrate that serum from HAdV-26 does not neutralize our HAdV-20-42-42 and vice versa.

To summarize, our studies into the HAdV-20-42-42 chimera demonstrate that we have identified a promising novel adenoviral vector to pursue both gene therapy and vaccine applications and therefore warrant further preclinical and clinical studies utilizing this vector.

## MATERIALS AND METHODS

### Origin and sequencing of HAdV-20-42-42

Strain 212 was isolated in the Skåne University Hospital, Lund, Sweden in 1978 (9). To obtain the complete genome sequence, it was propagated on human alveolar epithelial cells (A549), and the intracellular viral DNA was purified from infected cells (30). The genome was sequenced using Ion Torrent next-generation sequencing at the Uppsala Genome Center of the National Genomics Infrastructure (SciLifeLab; Uppsala, Sweden). The resulting reads were normalized to a 60-times coverage using BBNorm from the BBTools suite. The normalized reads were assembled de novo using Mira v4.9.5_2 (31), and the original sequence reads were mapped to the resulting consensus sequence using the Geneious mapper at the highest sensitivity with five iterations in Geneious 9.1.8 (32). After mapping the sequence reads to the *de novo* assembly, the final read coverage minimum was 51, the mean was 1048.4, and the read coverage’s standard deviation was 274.5. The new consensus sequence was annotated based on HAdV reference strain genome annotations, using the Annotate & Predict function of Geneious. The annotations were checked manually and edited. The genome sequence was submitted to the NCBI Nucleotide database with accession number MW694832.

### Phylogenetic analysis

Phylogenetic analyses were conducted based on the complete genome sequence and derived amino acid sequences of the entire hexon and penton base; and also on hexon loop 1 (delimited according to Yuan et al. (33)) and the fiber knob. For phylogenetic tree inference, multiple alignments were conducted using MAFFT (34), and phylogenetic calculations were performed using RAxML-NG v0.9.0 (35) based on alignments edited in trimAl v1.2 (36). Evolutionary model selection was performed using ModelTest-NG v0.1.5 (37). The robustness of the trees was determined with a non-parametric bootstrap calculation using 1,000 repeats. Phylogenetic trees were visualized using MEGA 7 (38), trees were rooted on the midpoint, and bootstrap values are given as percentages if they reached 75%. Recombination events were analyzed using SimPlot v3.5.1 (39).

### HAdV seroneutralization

Serological inhibition of HAdV-20-42-42, HAdV-35 and HAdV-5 transduction was evaluated over a collection of 103 serum samples from a cohort of healthy >50-year-old US citizens. Seroneutralization assays were performed using the protocol described in Sprangers *et al*. (25). Briefly, serum samples were heat-inactivated at 56 °C for 60 min, and then twofold dilutions were performed in a 96-well tissue culture plate. The dilutions covered a range from 1/16 to 1/4 096 in an end volume of 50 µl DMEM. Negative controls consisted of DMEM alone. After addition of 50 µl virus solution (1×10^8^ VP ml^−^ 1) to each well, a cell suspension (10^4^ A549 cells) was added to the well to a final volume of 200 µl. Following 24 h incubation, the luciferase activity in the cells was measured using a Steady-Glo luciferase reagent system (Promega). The neutralization titers were defined as the maximum serum dilution that neutralized 90 % of luciferase activity.

### Cell lines

HEK293 (human embryonic kidney cells: ATCC CRL-1573) were grown in Dulbecco’s modified Eagle’s medium (DMEM) supplemented with 10 % foetal bovine serum (FBS; Gibco, UK). A549 (human lung epithelial carcinoma: ATCC CCL-185) cells were grown in RPMI-1640, supplemented with 10 % FBS, 1 % penicillin/streptomycin (P/S), 1 % L-glutamine (Gibco) and 1 % Na-Pyr (Sigma, UK). Chinese hamster ovary (CHO) cells (40), CHO-CAR (40), various CHO-CD46 cells (41) were grown as described before. TC1-DSG2 cells, a kind gift of dr. A Lieber (University of Washington, Seattle, WA, USA), were grown in RPMI supplemented with 10% FBS and 20 mM HEPES.

### Construction of replication-incompetent recombinant HAd-20-42-42 vectors

The wild-type HAd-20-42-42 virus was plaque purified and propagated on HEK293 cells and purified by cesium chloride (CsCl) density gradient centrifugation. From the purified virus material full genomic DNA was isolated which served as starting material for the construction of the HAdV-20-42-42 plasmids.

### HAdV-20-42-42 cloning system

The HAdV-20-42-42 vector construction strategy was based on a three-plasmid system with sufficient homology between each of the plasmids to enable homologous recombination *in vitro* following the co-transfection in HAdV-5 E1-complementing HEK293 cells. Adapter plasmids that contain the left end of the HAdV-20-42-42 genome with E1 deletions and include either luciferase, EGFP or LacZ reporter genes, were generated first. Briefly, the adapter plasmid pAdApt20-42-42 (nt 1–461 of WT HAdV-20-42-42) contained the left inverted terminal repeat (ITR) and included an expression cassette consisting of a CMV promoter followed by a multiple cloning site, encompassing Luc, GFP or LacZ, and the SV40 poly(A) signal. This plasmid also contained the wild-type HAdV-20-42-42 nucleotides (nt) 3361-5908 to allow homologous recombination in HEK293 cells, with the intermediate plasmid carrying the HAdV-20-42-42 genome from IVa2 to L3 genes (nt 2088 to 18494). The right-end plasmid contained the HAdV-20-42-42 genome from L3 gene to the right ITR (nt 15373 to 35187) and had a deletion for the E3 region, while the HAdV-5 E4-ORF6 replaced the HAdV-20-42-42 E4-ORF6. The E3 region was deleted by PCR and standard cloning techniques exploiting a natural AscI site in the viral genome. To replace the native ORF6/7 by the homologue region of HAdV-5, three fragments were amplified by PCR. Two were designed to cover the region upstream and downstream from the ORF6/7 to be replaced. A third PCR was performed to obtain the HAdV-5 ORF6/7 (HAdV-5 GenBank accession no. M73260 nt 32963–34077) and partly overlapping with the other two PCR products. These three PCR products were then subjected to fusion PCR and cloned into the plasmid backbone to obtain the final right-end plasmid.

### Generation and production of HAdV-20-42-42-based adenoviral vectors

Adenoviral vectors HAdV-20-42-42-LacZ, HAdV-20-42-42-Luc and HAdV-20-42-42-GFP were generated by co-transfection of E1-complementing HEK293 cells with the adaptor plasmid, intermediate plasmid and the right-end plasmid. Prior to transfection into HEK293 cells, the three plasmids were digested with PacI to release the respective adenoviral vector genome fragments. The transfections were performed using Lipofectamine transfection reagent (Invitrogen; Carlsbad, CA) according to the manufacturer’s instructions. After harvesting of the viral rescue transfections, the viruses were further amplified by several successive infection rounds on HEK293 cell cultures. The viruses were purified from crude viral harvests using a two-step cesium chloride (CsCl) density gradient ultracentrifugation procedure as described before (14). Viral particle (VP) titers were measured by a spectrophotometry-based procedure described previously (42).

### Viral transduction assays

The transduction assay was performed on 96 well tissue culture plates (Costar). HSVEC were seeded at 1×10^4^ cells/well. The following day monolayers were washed with PBS and viral vectors, either encoding the luciferase or *β*-galactosidase gene, were added at the indicated VP/cell concentrations in their corresponding media without serum. In the experiments in which blood coagulation factor FX was used, this was pre-incubated with the vector for 30 min at 37 °C prior to addition to the cells. The FX coagulation factor was purchased from Cambridge Bioscience and used at a physiological concentration of 10 µg/ml. After 3 h incubation at 37 °C, the cells were washed and fresh medium supplemented with 10 % FBS was added, after which the cells were incubated for 48 h. For the readout of *β*-galactosidase gene expression, staining of the cells was performed using Galacto-Light Plus Assay Kit (Thermo Fisher Scientific). For the readout of luciferase activity, the plates were washed with PBS and the cells were lysed with 1× reporter lysis buffer (100 µl per well; Promega). Following a freeze-thaw cycle, luciferase (Luciferase assay system, Promega) measurements were performed with 20 µl of lysed cells in white opaque plates (Greiner BioOne) following the manufacturer’s instructions. The BCA protein quantitation assays (Thermo Fisher Scientific) were performed with 20 µl lysed cells. The results were recorded in a Victor X multilabel plate reader (Perkin-Elmer). The transduction level was expressed as luminescence relative light units per mg of protein per well (RLU/mg). All of the assays were performed with four replicates of samples.

### Receptor usage assays

Studies of receptor usage were performed using CHO cells, which were expressing (positive) or lacking (negative) for receptors of interest. The cells (CHO-CAR, CHO-sialic acid, CHO-DSG and CHO-CD46 (K1, BC1, BC2, C1 and C2) were seeded as four replicates in 96-well tissue culture plates and infected with HAdV-20-42-42-Luc or HAd5-Luc at a concentration of 1×10^4^ VP /cell. Cell cultures were incubated 3 h at 37 °C with 5 % CO2. Luciferase levels were measured at 48-72 hours post-infection with Victor X Multilabel plate reader (Perkin Elmer) following the manufacturer’s instructions. Luciferase transgene expression was presented as luminescence relative light units (RLU) per mg of protein. All assays were performed with four sample replicates for each experimental condition.

### *In vivo* biodistribution

All animal experiments were fully approved by University of Edinburgh Animal Procedures and ethics committee and performed under the UK Home Office license in accordance with the UK Home Office guidelines. Immunocompetent outbred MF1 male mice (Charles Rivers Laboratories) aged 8-10 weeks were used. The animals were organized in six groups with five animals in each group, except the control (PBS) groups which had 3 animals. In order to deplete circulating macrophages and more efficiently evaluate the transit of the virus at the whole organism level, 200 µl of clodronate liposomes (CL) was intravenously (i.v.) administered to corresponding groups at 48 h prior to virus administration.

Treatment groups were i.v. infected with a single dose (1×10^11^ VP diluted in 100 µl PBS) of HAdV-20-42-42-Luc or HAdV-5-Luc. Matched control groups were injected with 100 µl of PBS. At 48 hours post virus delivery, luciferase activity was imaged with the method IVIS Spectrum (CaliperLife Science, UK). Prior to the imaging procedure, 0.5 ml luciferin was injected to the mice. Animals were maintained under inhalational anesthesia (AB-G). Luciferase activity detected ranged from low (shown in blue) to high (shown in red) levels. After the imaging was completed, animals were sacrificed and their organs (liver, heart, spleen, kidney, intestine, pancreas and lungs) were collected for the quantification of the vector genomes as described before (13).

### Mouse immunization study

All animal experimentation was performed according to Dutch law and the Guidelines on the Protection of Experimental Animals published by the Council of the European Committee. Six-to-eight-week-old specific pathogen-free female Balb/c mice were purchased from Charles River and kept at the institutional animal facility under specified pathogen-free conditions. For prime immunization studies mice were immunized intramuscularly with HAdV-20-4-42 or HAdV-26 vectors (1×10^9^ VP or 1×10^10^ VP per mouse) expressing Luc. Two weeks post-immunization, mice were sacrificed and the induction of Luc-specific IFNγ-producing cells was measured by IFNγ ELISPOT. In brief, mouse splenocytes were stimulated with 15-mer peptide pool spanning Luc (luc pool), medium (control negative) or phorbol myristate acetate (PMA; positive control).

Mice antisera against HAdV-20-42-42 and HAdV-26 were cross-tested against both vectors in an adenovirus neutralization assay. Starting from a 1:16 dilution, and the sera were 2-fold serially diluted, as described elsewhere (25).

### Statistical analysis

Statistical analysis was performed with GraphPad Prism software. One-way ANOVA with the two tailed Student’s t-test was used for statistical parameter comparison between different groups. The parameters of significance are indicated in each figure caption: \**P*<0.05, \*\**P*<0.005, \*\*\**P*<0.001 and NS, not statistically significant, *P*>0.05. The presented *in vitro* results are averaged data from at least three different experiments with four experimental replicates per condition. The *in vivo* experiments were performed with a minimum of five animals per group. Errors bars represent the standard error of the mean (SEM). Statistical analyses were performed with SAS version 9.4 for Figure 6. Non-inferiority testing across dose was performed on log10-transformed data, with HAdV-26-Luc as a reference and a pre-specified margin of 0.5 log10.

## ACKNOWLEDGMENTS

This study was made possible by funding from FP7 Marie Curie Actions via the ADVEC consortium (grant agreement no.: 324325). This project has received funding from the European Union’s Horizon 2020 research and innovation programme under grant agreement number 825670. The research of GLK and TP is supported by the National Research, Development and Innovation Office (OTKA NN128309), and that of GLK also by the János Bolyai Research Scholarship of the Hungarian Academy of Sciences. The assembly and phylogenetic calculations were performed using resources provided by KIFÜ, Hungary.

